# Mechanisms of speed encoding in the human middle temporal cortex measured by 7T fMRI

**DOI:** 10.1101/2021.02.01.429141

**Authors:** Anna Gaglianese, Alessio Fracasso, Francisco G. Fernandes, Ben Harvey, Serge O. Dumoulin, Natalia Petridou

## Abstract

Perception of dynamic scenes in our environment results from the evaluation of visual features such as the fundamental spatial and temporal frequency components of a moving object. The ratio between these two components represents the object’s speed of motion. The human middle temporal cortex hMT+ has a crucial biological role in the direct encoding of object speed. However, the link between hMT+ speed encoding and the spatiotemporal frequency components of a moving object is still under explored. Here, we recorded high resolution 7T blood oxygen level-dependent BOLD responses to different visual motion stimuli as a function of their fundamental spatial and temporal frequency components. We fitted each hMT+ BOLD response with a 2D Gaussian model allowing for two different speed encoding mechanisms: 1) distinct and independent selectivity for the spatial and temporal frequencies of the visual motion stimuli 2) pure tuning for the speed of motion. We show that both mechanisms occur but in different neuronal groups within hMT+, with the largest subregion of the complex showing separable tuning for the spatial and temporal frequency of the visual stimuli. Both mechanisms were highly reproducible within participants, reconciling single cell recordings from MT in animals that have showed both encoding mechanisms. Our findings confirm that a more complex process is involved in the perception of speed than initially thought and suggest that hMT+ plays a primary role in the evaluation of the spatial features of the moving visual input.

## Introduction

Encoding of visual features from dynamic visual images is essential in humans and nonhuman primates to reconstruct the visual scene and rapidly respond to the ever changing environment. Among visual areas, the human homologue of the macaque middle temporal cortex (hMT+ also known as V5) has been shown to play a functional role in the encoding of features such as the spatial and temporal frequency components of visual motion stimuli^1–4^. Using electrocorticography, we recently showed that hMT+ neuronal populations separated visual motion into its spatial and temporal components, with speed preferences changing in accordance with the fundamental spatial frequency of the visual stimuli, rather than being tuned for a particular speed of the attended moving stimuli ^5^. These findings, paired with single cell recording studies in animals, describe hMT+ neurons as spatiotemporal frequency sensors for motion extraction ^3,4,6^. However, debate continues about the speed encoding mechanisms of MT since pure speed tuning encoding has been reported in different MT cells in primates ^7–10^. One major issue of animal single-cell recordings and human electrocorticography measurements is the reduced coverage of MT+ due to the closely-spaced recording sites. Therefore, it remains elusive whether there is a functional organization within the complex for the different mechanisms of speed encoding, i.e. separable tuning for spatial and temporal frequencies vs pure speed tuning. The rapid development of ultra-high field (7 Tesla, 7T) functional Magnetic Resonance Imaging (fMRI) allows us to reveal the fine-scale functional organization of the human cortex in vivo ^11,12^. Many 7T fMRI studies have been carried out in primary visual cortex V1, although studies have been recently extended to reveal the fine-scale functional organization of the human extrastriate cortex and association areas ^13–21^. A recent high spatial resolution 7T fMRI study in hMT+ in particular, has demonstrated a functional organization into columnar clusters with preferences for horizontal or vertical motion, similar to the columnar organization in monkeys ^22^. However, human research to date has tended to focus on the spatial organisation of responses to the location and direction of motion in hMT+, rather than the mechanisms involved in the encoding of speed of motion.

Here, we disentangle whether and to what extent the hMT complex is directly tuned for the speed of motion or encodes speed via tuning to fundamental spatiotemporal properties of the visual motion stimuli. We used high resolution 7T fMRI to characterize the organization of hMT+ Blood Oxygenation Level Dependent (BOLD) response amplitudes for different combinations of fundamental spatial and temporal frequency components of visual motion stimuli. We modelled BOLD responses with a 2D Gaussian model that allows for either speed tuning encoding or separable tuning of the spatial and temporal frequencies components. We were able to characterize the mechanisms involved in the encoding of speed of motion and to demonstrate that both encoding mechanisms occur in hMT+, with the majority of the complex exhibiting temporal and spatial frequency selectivity.

## Methods

Five healthy volunteers (all male, mean age + SD = 36.2 ± 3 years) participated in the study after giving written informed consent. The study was approved by the Ethics Committee of the University Medical Center of Utrecht in accordance with the Declaration of Helsinki (2013) and the Dutch Medical Research Involving Human Subjects Act.

### hMT+ localizer stimulus

Area hMT+ was functionally identified based on responses to moving compared to stationary visual stimuli, as conventionally used in literature ^23,24^. We used a full field high-contrast square-wave black-and-white dartboard patterns instead of standard random dots to match the contrast of the visual stimulus used in the visual motion stimulation experiment. During the motion condition, the dartboard pattern expanded from the fixation point for 10s with a temporal frequency of 5 Hz, interleaved with a stationary period of 10s during which the same dartboard was presented static. The stimuli subtended a visual angle of 30.7×16.1°.

### Visual motion stimulation

The visual motion stimulation consisted of five runs of high-contrast square-wave black and white dartboard patterns presented with three different fundamental spatial and temporal frequency combinations (0.33 cycle/deg;1Hz, 0.33 cycle/deg; 3Hz, 0.33 cycle/deg; 5Hz, 0.2 cycle/deg; 3Hz, 1 cycle/deg; 3Hz). The range of spatial frequencies was based on our previous EcOG study and fMRI recordings in humans showing peak responses in hMT+/V5 around 0.33 cycle/deg and reaching the minimum response amplitude for a spatial frequency of 1.24 cycle/deg ^25–29^. Given that the speed of motion of each square-wave dartboard presented is defined by the ratio of temporal to spatial frequencies, speeds of 3deg/sec, 9 deg/sec, and 15deg/sec were presented respectively. Each run is either classified as a fast (15 deg/sec), intermediate (9 deg/sec) or slow (3 deg/sec) moving stimuli, depending on the fundamental spatial and temporal frequency that gives origin to the stimuli speed. The fast and the slow speed respectively were presented twice by using two different spatiotemporal frequency combinations of the moving dartboards (see fig. 1).In each run, we presented only one spatiotemporal frequency combination for a total of 26 trials. Each run lasted 5min 10sec. The dartboard pattern expanded from the fixation point for 1s alternating with a baseline period during which the dartboard was static to enhance the effect of motion. Baseline periods were of variable length ranging from 6s to 15s, presented in a pseudo-randomized order. Three additional baseline periods of 24s were randomly added to allow the BOLD response to return to baseline. To maintain fixation and consistent level of arousal, participants were instructed to press a button when the central fixation dot changed color from red to green and viceversa (Fig1). Participant performance was recorded via Matlab software, and response accuracy was consistently above chance for each run and participant.

**Figure 1:**
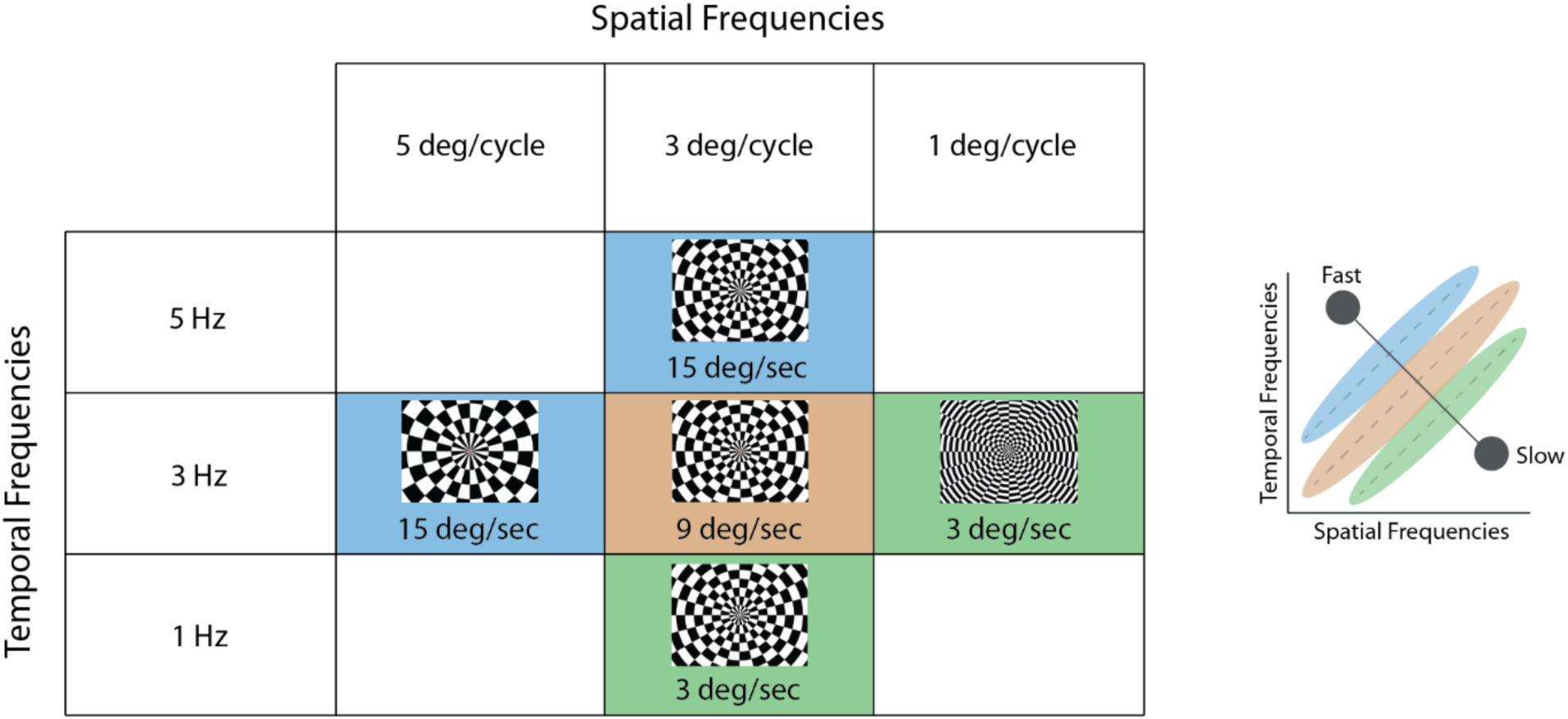
Graphical summary of the visual motion stimuli (high-contrast black-and-white dartboard) presented during the five runs of the study following the localizer run. Each run is either classified as a fast (represented by blue background color), intermediate (represented by orange background colour) or slow (represented by green background colour) moving stimuli, depending on the spatial and temporal frequency that gives origin to the stimulus speed. Graphic on the right provides a simplified depiction of the stimulus space.

## fMRI methods

### fMRI data acquisition

Functional MRI data were acquired using a Philips 7T scanner equipped with a volume transmit (Nova Medical, USA) and two high-density16-channel surface coils ^30^. The surface coils covered the left and right lateral occipital pole of the participant to maximize the signal-to-noise (SNR) and BOLD sensitivity in the left and right hMT+. A gradient echo echo-planar-imaging (EPI) sequence was used for both the localizer and the visual motion stimulation experiments. Functional images for the localizer were acquired every 1.8s, with an echo time (TE) of 27ms, an isotropic voxel of 1.5mm and 27 coronal slices covering hMT+ bilaterally. For the visual motion stimulation experiment we acquired 15 coronal slices at a fast temporal resolution of 0.849ms, TE=27ms, and an isotropic voxel resolution of 1.4mm. For both functional acquisitions, EPIs were acquired with a SENSE factor of 2 in the right-left direction. High-resolution T1-weighted anatomical MRI images were acquired with a 32-channel head coil (Nova Medical, MA, USA) in a different session at a resolution of 0.8×0.8×0.8 mm. Repetition time (TR) was 7 ms, TE was 2.84 ms, and flip angle was 8 degrees. Visual stimuli were projected onto a 27 × 9.5 cm screen placed inside the magnet bore behind the participant’s head, using a projector (Benq W6000, 1600×538 pixels display resolution). Participants viewed the projected visual stimulus via a mirror and prism glasses, at an effective distance of 41cm.

### Data pre-processing

All the pre-processing steps were performed using AFNI (Analysis of Functional NeuroImages, https://afni.nimh.nih.gov/). First, the functional data of the localizer and visual motion stimulation runs were corrected for motion and aligned to the first image of the first run of each session respectively using the function 3dVolreg. Subsequently, low frequency signal intensity drifts were removed with quadratic detrending using the 3dDetrend function. No spatial smoothing was employed. The visual motion stimulation runs were non linearly co-registered to the localizer using the 3WarpDrive function. To avoid time series interpolation of the visual stimulation runs, we extracted the Regions of Interest (ROIs) in the localizer space (see Localization of hMT+) and co-registered them to the visual motion stimulation runs using the inverse of the obtained transformation matrix.

The T1-w anatomical images were segmented automatically using the MIPAV software package implemented in CBStool (https://www.nitrc.org/projects/cbs-tools/). White matter and pial surfaces were generated and then imported in SUMA (afni.nimh.nih.gov). For each participant, maps obtained from the functional runs (see below) were co-registered on the pial surface using first the function 3dAllineate with mutual information as cost function, and then non-linearly using the 3dWarpdrive function.

### Localization of hMT+

For each participant, left and right hMT+ areas were functionally defined from the localizer runs by contrasting responses for the moving and stationary high contrast black and white dartboard stimuli. All statistical computations were performed at a single participant level using a general linear model (GLM) with a standard gamma variate hemodynamic response function approach, using the 3dDeconvolve function in AFNI. For each run, outliers due to residual motion were detected via 3dTOutCount function and included in the GLM analysis as regressors of no interest. Voxels that exhibited significant responses for moving vs stationary dartboards (p<0.001, Bonferroni corrected) and located within the left and right hMT+ anatomical landmarks ^31^ on the EPI space were selected to define hMT+ ROIs.

### Quantification of BOLD responses for the visual motion stimulation

The BOLD responses to each combination of spatial and temporal frequencies presented during the visual motion stimulation runs were estimated for all voxels using a finite impulse response deconvolution approach ^32–34^ implemented in mrTools (available for free download at http://gru.stanford.edu/doku.php/mrTools/overview), a software package running in MATLAB. The response to a given stimulus type was quantified for each voxel by the amplitude of the BOLD response. Only amplitude values within each hMT+ ROI were selected for further analysis. We computed the significance of the effect of the spatial and temporal frequency components of the visual motion stimulation on the hMT+ BOLD response amplitudes by two-way ANOVA within participants. Furthermore, to investigate if the BOLD response amplitude changes according to the spatiotemporal frequency combination of the stimuli rather than the speed per se, we performed a two-sample t-test between the 3deg1Hz and 1deg 3Hz spatiotemporal frequency combination (both representing 3deg/sec speed of motion of the visual stimuli) and 3deg5Hz and 5deg3Hz (both representing 15deg/sec speed of motion).

### Tuning Model of hMT+ BOLD responses

We ask whether the BOLD responses in hMT+ are consistent with tuning for the fundamental spatiotemporal frequency combination of the presented moving visual stimuli or whether they are consistent with pure speed tuning. To answer this, we compared the measured BOLD response amplitudes with the predicted BOLD response amplitudes obtained by both a spatiotemporal frequency tuning model and a speed tuning model ^4–6,35^. The first model retains separate and independent responses for the spatial and temporal frequency components of the visual stimuli. The second model describes direct encoding of the speed of motion, resulting in a preference for the same speed at different spatial frequencies, with temporal frequency tuning varying in accordance with the spatial frequency. Both models are represented by a two-dimensional Gaussian function with the addition of an extra parameter Q which allows to characterise the two different types of tuning. A value of Q equal to zero (Q=0) describes separable responses for each spatial and temporal frequency combination of the stimuli. A value of Q equal to 1 (Q=1) describes tuning for particular speeds, i.e., predicts the same optimal speed at different spatial frequencies. Both models are described by the equation below:

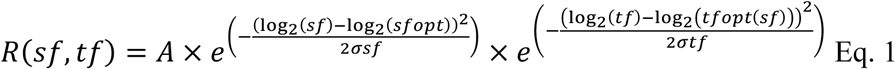

Wherelog_2_(tfopt(sf)) is defined as:

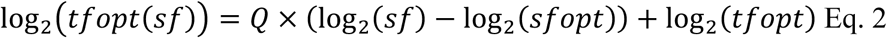

The term A is the peak of the BOLD response amplitude, sfopt and tfopt are the optimal spatial and temporal frequency, and σsf and σtf are the bandwidths of the spatial and temporal frequencies tuning curves. We fitted the BOLD response amplitudes of each voxel in each hMT+ ROI to both the spatiotemporal frequency and the speed tuned models (setting Q = 0 or Q = 1 respectively). For each voxel and model, three parameters were estimated: a) optimal fundamental spatial frequency b) optimal fundamental temporal frequency c) variance explained. We assigned for each voxel of each hMT+ ROI an optimal tuning model (separable spatiotemporal frequency tuning or speed tuning) based on the best fit (higher variance explained to one of the two models).

### Cross validation

We cross-validated each model’s goodness of fit in predicting the BOLD response amplitude for each voxel by fitting the BOLD response amplitudes of one half of the measured data and testing how well the resultant parameters predict the BOLD response amplitudes in the complementary half, assessed by the variance explained (Mante et al., 2005). For this purpose, two independent halves of the data are needed. Since in our visual motion stimulation paradigm we presented each spatiotemporal condition in a unique run we split each run in two halves according to the incidence of the second 24s baseline period. Hence, to minimize the possible effect of the BOLD response of the last trial of the first half split of the run on the first trial of the second half split. We applied this approach for both the speed tuning model (parameter Q = 1) and the spatiotemporal frequency tuning model (Q=0). Voxels with variance explained below 0.1 in both models were discarded from the subsequent analysis. On average 39.2±4.27 percent of voxels were discarded from each bilateral hMT+ ROI. A two-sided paired t-test between the average variance explained of each model within each bilateral hMT ROI+ was computed to define the model that best represented the measured BOLD responses. To test the reproducibility of the model fitting, the same analysis was performed by fitting the BOLD responses on the second half of the data and testing on the complementary first half of the data for both Q=0 and Q=1 model. We define the repeatability of the best model fits for each ROI using a permutation test (n = 1000 shuffle repetitions) on each bilateral hMT+ ROIs variance explained obtained from the two split halves for the two models.

### Separable tuning of hMT+ BOLD responses for temporal and spatial frequencies

For each half split and each ROI we obtained a spatial map of optimal spatial and temporal frequencies. We quantified the reproducibility of these maps by computing a Spearman correlation coefficient between the two-half split spatial distribution of the sfopt and tfopt parameters. Based on our previous ECoG study in humans using the same paradigm ^5^ and neurophysiological findings in animals ^25,26,36^, we expected hMT+ to be more tuned toward low spatial frequency rather than high spatial frequency. To quantify this effect, we classified the distribution of the optimal fundamental spatial frequencies obtained from the Q=0 model in k clusters. To guide the choice of the number k of clusters to be used to classify the spatial frequencies we computed the within sum of squares accounting for the number of clusters, by using the Bayesian information criterion BIC. We then classified independently the spatial frequency parameters using k-means. Mean and standard deviation of the center of clusters and cluster size for each hMT+ ROI were computed across participants.

## Results

### hMT+ BOLD responses showed differential responses for each combination of spatial and temporal frequency of the visual motion stimuli

Regions of interest (ROIs) for hMT+ were defined for each hemisphere based on the functional localizer. The mean grey matter area for the right and left ROIs were 451±90.5 mm^3^ and 251±18.4 mm^3^ respectively.

Figure 2 depicts the ROI extent in the anatomical space. Each voxel within each ROI was assigned a spatiotemporal frequency combination preference in accordance with the maximum BOLD response amplitude across combinations.

**Figure 2:**
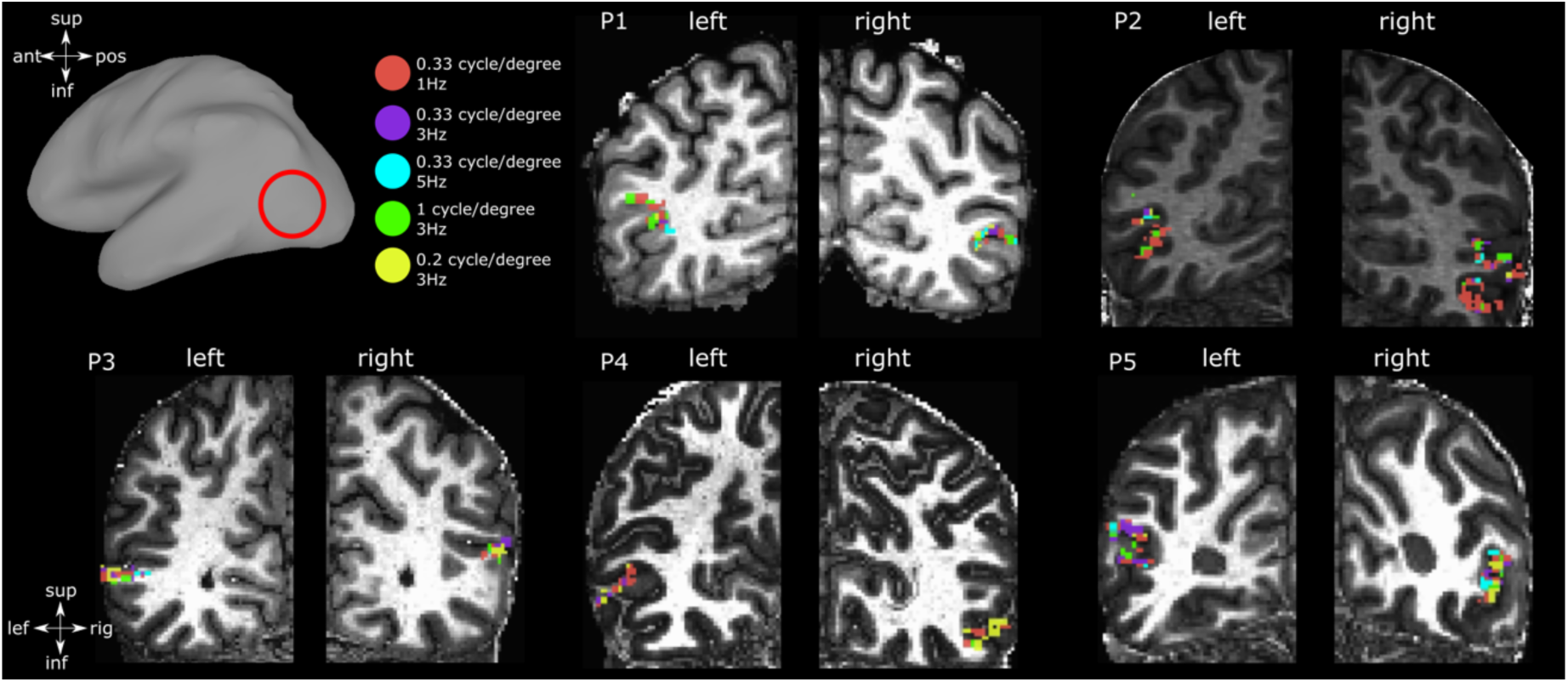
Spatiotemporal frequency preference within hMT+ ROI of each participant plotted on each anatomical space. The amplitude with the highest value across all spatiotemporal frequency conditions determined the preferred spatiotemporal frequency combination of each voxel.

The effect of each fundamental spatiotemporal frequency combination on the bilateral hMT+ ROIs was significant across participants (one-way ANOVA, F = 4.45, p = 0.0005). The amplitude of BOLD responses was significantly different for each measured spatial and temporal combination. Four out of five participants exhibited significantly different BOLD response amplitudes (p<0.05) for the two pairs of spatiotemporal frequency combinations leading to the same speed of motion of the presented dartboard (3deg1Hz - 1Hz3deg and 3deg5Hz - 5deg3Hz respectively).

### hMT+ BOLD response amplitudes were mainly characterized by independent tuning for spatial and temporal frequency

The two-dimensional Gaussian models allowing independent tuning for spatial and temporal frequency (Q=0) or tuning dependent on speed (Q=1), were able to characterize the hMT+ BOLD amplitude responses (Figure 3a). For each model we fitted each voxel’s BOLD response amplitude for each combination of the fundamental spatial and temporal frequency on one split half of the data and computed the variance explained by the resulting model in the second complementary half. Overall, for each participant’s bilateral hMT+ ROIs, in cross validation, the Q = 0 model explained significantly more variance than the speed encoding model Q = 1 (two-sided t-test, *p*<0.001 for each participant, see Figure 3a). Each bilateral hMT+ ROIs also included a number of voxels exhibiting higher variance explained for the Q=1 model compared to the Q=0 model. The percentage of grey matter area within each bilateral hMT+ ROI of each participant that is better explained by the Q=0 or Q=1 model respectively is shown in Figure 3b. These two groups of voxels exhibiting different speed tuning profile are repeatable across the split run validation (fig. 3c and d). The group of Q=1 voxels is highly repeatable (p<0.001) across subjects, although it does not reach significance in every participant.

**Figure 3:**
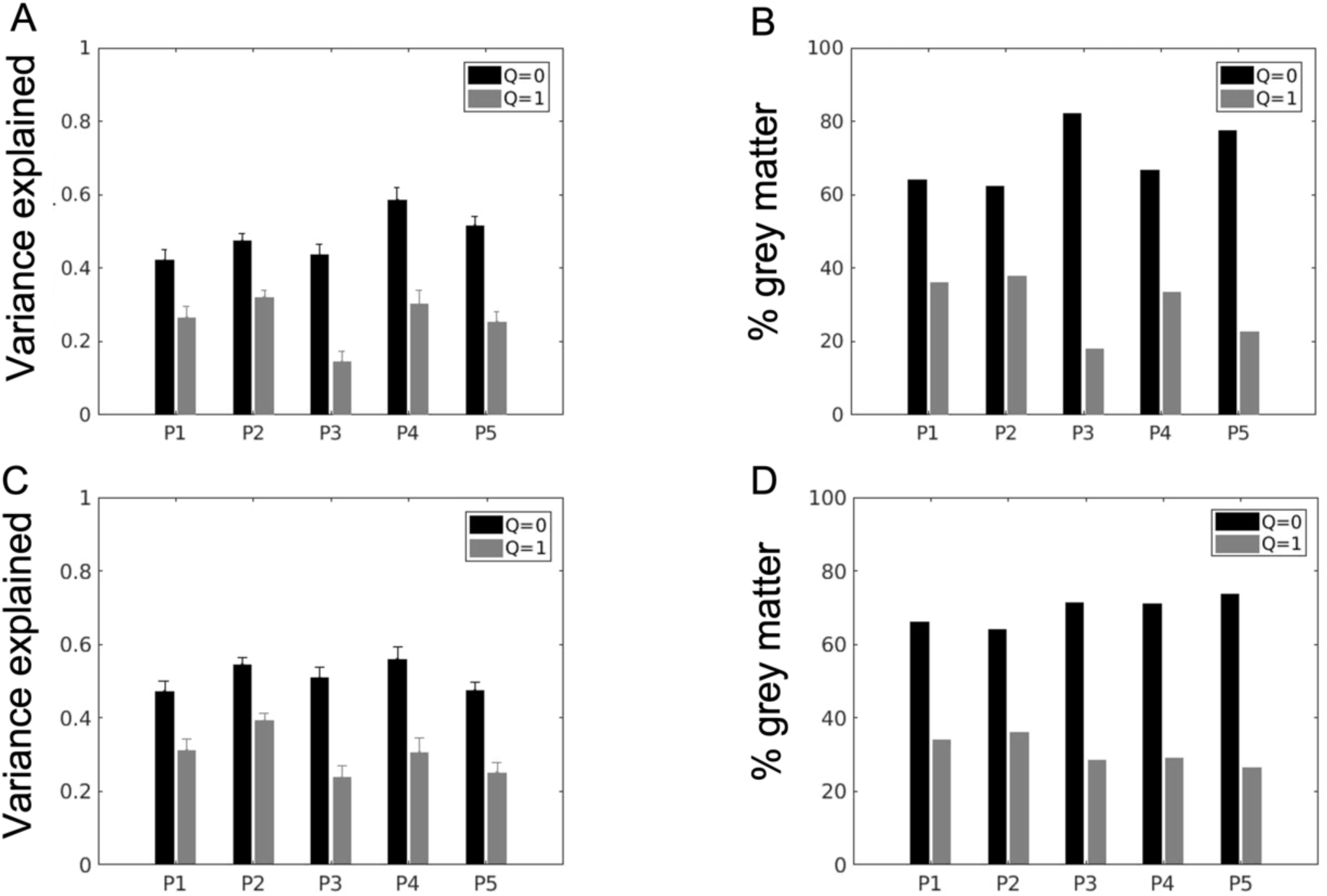
A) Single participant variance explained in split-run cross validation (fitted on first half of the run and validated on second half) by Gaussian tuning models with independent spatial and temporal frequency tuning (Q=0) and tuning for speed (Q=1). Each bar represents the mean variance explained and standard error for bilateral hMT+ ROI voxels of each participant (P1 to P5, x-axis). B) Single participant percentage of voxels within hMT+ ROIs exhibiting separable spatiotemporal frequency tuning (Q=0, black bars) or speed tuning, i.e. same temporal frequency preference for the different spatial frequency of the moving dartboard (Q=1, grey bars) C) D) Same as A and B tested in split-run cross validation on second half and validated on first half.

Figure 4 shows the histograms of the optimal fundamental spatial and temporal frequencies, variance explained, and speed preferences within bilateral hMT+ ROI voxels of each participant for the spatiotemporal frequency tuning model Q=0. Median optimal fundamental spatial and temporal frequency values were consistent across participants.

**Figure 4:**
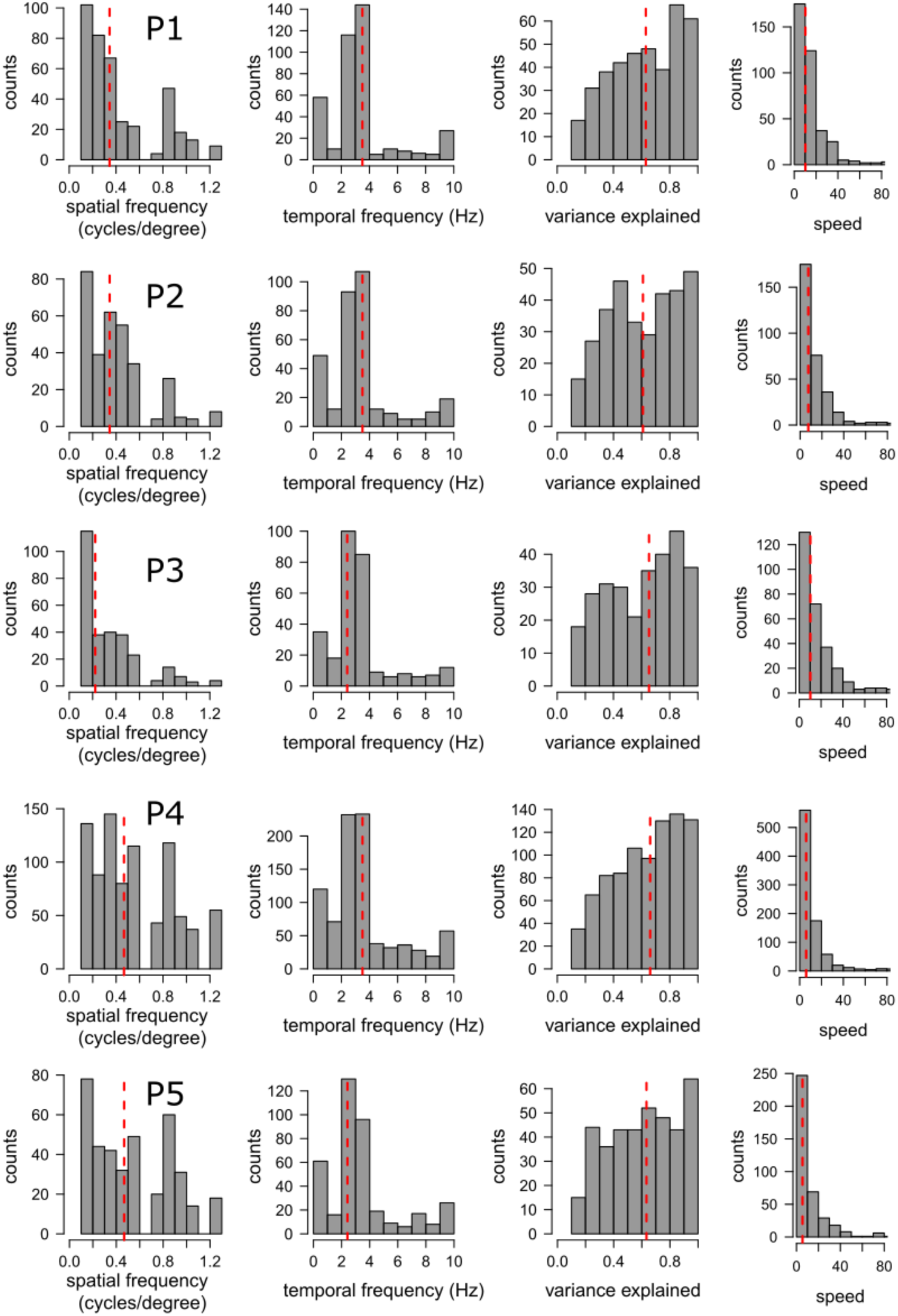
Histograms of estimated optimal temporal frequencies, spatial frequencies, variance explained and speeds obtained in cross validation for each participant’s bilateral hMT+ ROI. Red dashed line shows the median values across voxels.

We further tested the reproducibility of the cortical organisation of the optimal spatial and temporal frequency preferences across the two half splits of the data using Spearman’s correlation. Only maps of the spatial frequency preferences exhibited a significant correlation in all the participants’ bilateral hMT+ ROIs (r2 = 0.64, 0.59, 0.79, 0.65, 0.61 respectively, p<0.0001). Optimal fundamental spatial frequency values for each participant and each half split were then classified in two clusters respectively using kmeans. K = 2 was based on the optimum value displayed by the BIC score (fig. 5A). Optimal spatial frequency clusters for the complete run and each half split for a representative participant are shown in Figure 6A-C. A parallel-coordinates plot for spatial frequencies is shown in fig. 6D-E in which the starting point of each line on the left side of each plot indicates the spatial location of the voxel and the cluster classification, and the ending point the correspondent classification in the second half. Mean centroids across participants (Fig. 5B) were consistent and centered on 0.20±0.013 cycle/degree (low spatial frequencies cluster) and 0.79±0.035 (high spatial frequencies cluster). The percentage of voxels normalized by the size of each cluster is shown in Figure 5C.

**Figure 5:**
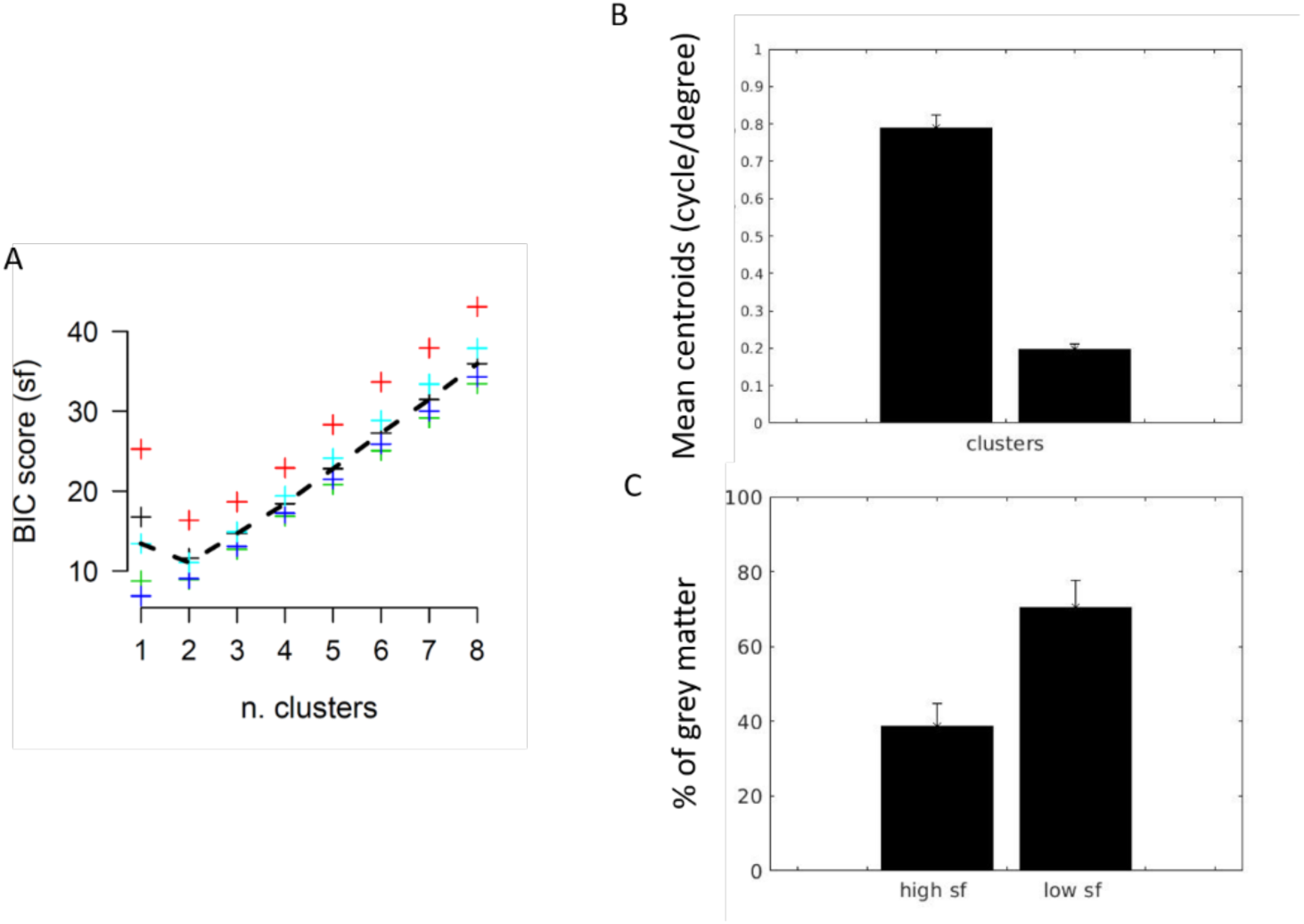
A) Bayesian Information Criterion score as a function of the number k of clusters for the optimal spatial frequency values. Each colour represents a single participant. Dashed line represents the mean score across participants. K = 2 was selected for the optimal number of clusters B) Mean centroids value and standard deviation across participants hMT+ ROIs for each spatial frequency cluster C) Mean percentage of grey matter and standard deviation across participants hMT+ ROIs for each spatial frequency cluster.

**Figure 6:**
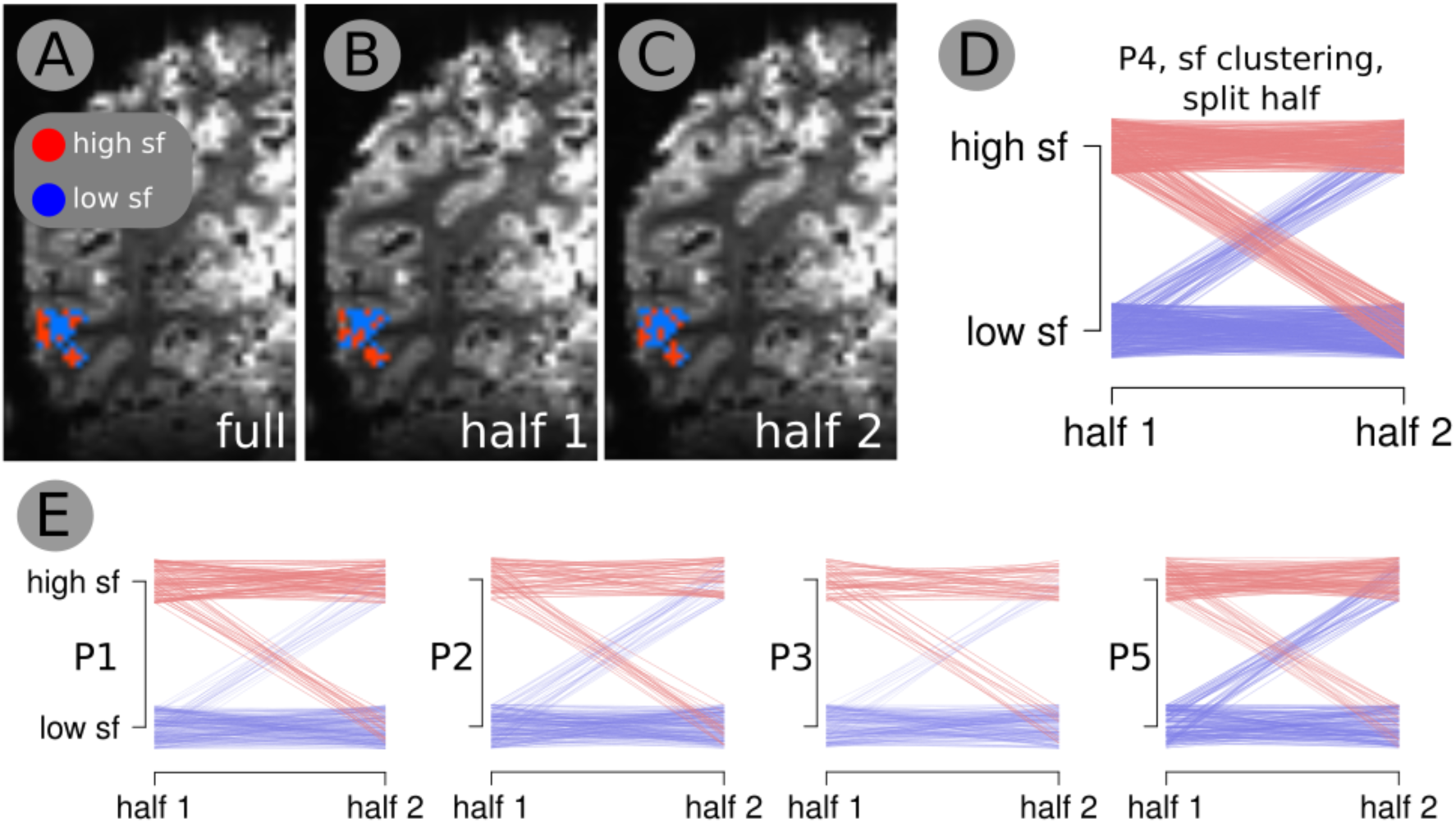
Optimal spatial frequency clusters and their reproducibility for a representative participant. A-B-C Spatial frequency clusters represented on the cortex for the entire run, half 1 and half 2 respectively. D) Parallel-coordinates plot. It represents optimal spatial frequency of each voxel in the first (left side) and second half (right side) run respectively. Y axis maps voxel coordinates in space. E) Same as D for each measured participant

## Discussion

In the current study we investigated the speed encoding mechanisms of hMT+ BOLD responses in response to different combinations of the fundamental spatial and temporal frequency components of visual motion stimuli. Overall, our results support the coexistence of two sub-populations within the complex exhibiting two different mechanisms of speed encoding. The bigger cluster exhibited separable tuning for the spatial and temporal frequency components of the visual motion stimuli, where the neuronal population speed preference changes in accordance with the spatial frequency. The smaller cluster exhibited direct speed tuning, where the same speed preference was maintained for different spatial frequency components of the stimuli.

These two different mechanisms have been previously reported in the literature. The first and more conventional one foresees direct speed tuning, where the same speed preference is maintained for different spatial frequency components of the stimuli. The second proposes independent tuning for the spatial and temporal frequency components of the visual stimuli, where the neuronal population speed preference changes in accordance with the spatial frequency. Animal studies using intracellular recordings ^3,4,6,35^ have shown in MT/V5 the presence of both mechanisms and in particular: 1) the existence of a percentage of MT cells responding to the speed of motion of the presented visual stimuli ^7,9^, 2) separable neuronal response within MT/V5 neuronal population ^1,2^ and 3) a continuum of both mechanisms ^37^. In humans we recently shown using electrocorticography ECoG that sampled neuronal population within the complex exhibited distinct and independent selectivity for spatial and temporal frequencies of the visual stimuli ^5^. Our results were limited to the coverage of the ECoG electrodes, possibly hiding smaller neuronal populations tuned for speed. We were now able to investigate the speed encoding mechanisms of the entire hMT+ complex with higher detail, bridging the gap between single cell recordings in animal studies and electrophysiological recordings of single neuronal populations in the human brain and reconciling the different results reported in the literature.

Moreover, for the regions of the complex exhibiting separable responses we further tested the spatial selectivity for spatial and temporal frequency by estimating the optimal fundamental spatial and temporal frequency values for each voxel of each participant’s hMT+ ROI. Optimal spatial frequency maps were highly reproducible within participants, in line with the hypothesis that visual areas with specific visual field maps such as MT/V5 exhibit specific responses for the spatial frequency component of the perceived stimuli. Indeed, tuning for spatial frequencies in the occipital cortex has been shown using optical imaging in cat and fMRI in humans ^25,26,36^, by showing a decrease in optimal spatial frequency tuning moving from V1 to V3 and to extrastriate cortex such as MT+. It has been shown ^26^, using fMRI in humans, low pass tuning responses for spatial frequency in V5/MT, exhibiting a significant drop in responses for spatial frequencies above 0.4 cycle/degree. In our dataset we measured the same effect: the optimal spatial frequencies were distributed along two clusters, peaking respectively on low spatial frequencies (0.20 cycle/degree) and on high spatial frequencies (0.79 cycle/degree), where the largest number of voxels in the entire hMT+ complex was tuned for the low spatial frequency cluster. Although the limited spatial frequency sampling of our experiment does not allow us to draw a firm conclusion, we suggest that this effect may reflect the change in eccentricity across the visual field maps, or different responses in the MT and MST (or TO1 and TO2) subdivisions of the complex. A single neuron recording study in the homologue V5 area in monkeys showed different spatial frequency preferences within the area (higher for MT and lower for MST) in accordance with the increase in eccentricity in MST compared to MT ^23^. In humans, the visual field map TO2 has larger pRFs than TO1 ^38,39^. Also, within both of these visual field maps, pRF sizes increase with eccentricity. Spatial frequency preferences typically decrease where pRF sizes increase, at higher eccentricities and in visual field maps with larger pRF sizes ^27^.

Finally, optimal temporal frequency tuning was not reproducible within participants. This can be due to the range of the temporal frequencies used in our experiment (from 1Hz up to 5Hz). It has been shown that the optimal contrast sensitivity of the primate visual system is found at approximately 8 Hz ^40–42^. A recent fMRI study in humans shows a peak at around 10Hz across visual areas independent of pRF size ^41^. Further studies exploring a wider range of spatial and temporal frequencies may help elucidating the spatial organization of the complex with higher detail.

Overall, our findings suggest that the majority of hMT+ responses to speed change in accordance with the spatial frequency component of the visual motion stimuli. We speculate that speed tuning properties may emerge from non-linear integration of patches within the MT complex preferring the same speed but different spatial frequency. Then, at a later stage, this information is computed in other subregions within the complex as suggested by the presence of small patches showing speed tuning properties rather than separable responses. Moreover, the fact that hMT+ exhibited the same properties as the primary visual cortex V1 in encoding basic features of a visual stimuli, such as the spatial and temporal frequency components, is consistent with previous studies in both humans and primates showing that area MT receives, and is able to process, fundamental properties of the visual input directly from the thalamus, bypassing the V1 ^43–45^ and could explain the absence of deficit in biological motion perception in patients affected by congenital visual deprivation ^46^. This fundamental low level mechanism of the hMT complex in processing visual motion features could explain the multisensory role of this area in encoding motion via other sensory modalities such touch and hearing ^47–50^. Indeed, asensory specific areas rely on the process of task information (e.g. motion) responses based on specific low-level properties of the input regardless the sensory modalities in which they are delivered ^51,52^.

## Conclusion

We provided evidence of the coexistence within hMT+ of a functional selectivity for spatial frequency, with speed preference changing in accordance with the fundamental spatial component of the presented visual motion stimuli and of a mechanism of pure tuning for the speed of the motion. These findings suggest that speed encoding in hMT+ is more complex than initially thought and underline the role of this area in computing feature properties of visual stimuli in a similar manner as primary visual cortex.

## Acknowledgment

This work was supported by the Netherlands Organization for Scientific Research (NWO), Vidi Grant number 13339 (N.P.) and by the National Institute Of Mental Health of the National Institutes of Health under Award Number R01MH111417. The content is solely the responsibility of the authors and does not necessarily represent the official views of the National Institutes of Health. A.F. is supported by a grant from the Biotechnology and Biology research council (BBSRC, grant number: BB/S006605/1) and the Bial Foundation, Bial Foundation Grants Programme 2020/21, A-29315;

## Data and code availability statements

All the analysis code can be obtained by emailing a request to the first author. Code used for the speed modelling results is available on a github repository: https://github.com/befrancislikeme/Encoding-spatiotemporal-frequencies-hMT. Sharing of data at this stage is not possible because we do not have consent from the participants to share their data. We will investigate if possible to get permission from the Ethics Review Board of the University Medical Center Utrecht to make anonymized processed data available upon request.

